# Effects and phenotypic consequences of transient thyrotoxicosis and hypothyroidism at different stages of zebrafish *Danio rerio* (Teleostei; Cyprinidae) skeleton development

**DOI:** 10.1101/2024.06.08.598073

**Authors:** Vasily Borisov, Fedor Shkil

## Abstract

Thyroid hormones (THs) are one of the main regulators of remodeling, homeostasis and development of skeletal tissues in teleosts, and the effects of hypo- and hyperthyroidism on skeleton are among the objectives of research in the fields of fishes development and evolution. However, in most experimental models used, the altered THs status is a constant characteristic of the developing organism, and the observed phenotypic outcomes are the cumulative consequences of multiple THs induced developmental changes. The effects of the transient fluctuations of THs content on the skeleton development have been studied much less. Here, we present experimental data on the developmental effects and phenotypic consequences of transient, pharmacologically induced thyrotoxicosis and hypothyroidism at different stages of zebrafish ossified skeleton patterning. In accordance with the results, skeleton structures differ in the timing and degree of THs sensitivity. Some of them displayed a notable shift in the developmental timing and rate, whereas other demonstrated a subtle or absence of reaction in respond to changes of THs content. The developmental stages also differ in THs sensitivity. A relatively short developmental period has been revealed, during which changes in THs level (mainly thyrotoxicosis) sharply increase the developmental instability and plasticity, leading to phenotypic consequences comparable to those in fish with permanently altered THs status. These findings allow us consider this period as a critical developmental window.

**Summary statement:** Study of the developmental effects and phenotypic consequences of acute transient changes in thyroid hormones content have identified a critical developmental window for zebrafish’s skeleton

**Ethics statement:** All procedures with fish were carried out according to the guidelines and following the laws and ethics of the Russian Federation, and approved by the ethics committee of the Severtsov Institute of Ecology and Evolution, Russian Academy of Sciences (Approval ID: N 95 issued on 27.05.2024).

## Introduction

Vertebrate skeleton development is a complex process regulated by a variety of factors, among which thyroid hormones (THs), products of hypothalamic-pituitary-thyroid axis (HPT axis), play a role that could not be overestimated. They have diverse effects on the remodeling, homeostasis and development of skeletal tissues, including the stimulation and regulation of coordinated developmental changes, such as the larvae-juvenile transition and metamorphosis (Bohnsack and Kahana, 2013, Kim and Mohan, 2013, McMenamin and Parichy, 2013, Holzer and Laudet, 2015, Deal and Volkoff, 2020, Keer et al., 2019, Keer et al., 2022, Roux et al., 2023).

The mechanism of action of THs is ancient and conservative in vertebrates (Norris and Carr, 2020). Thyroxine (T_4_) is a prohormone that is synthetized and released by the thyroid follicles, and converted in different tissues under the action of deiodinase enzymes into the active form of THs, triiodothyronine (T_3_). The latter modulates the developmental processes via binding with the transcriptional factors, thyroid hormone receptors, thereby determining the timing and rate of expression of target nuclear and mitochondrial genes, including genes involved in homeostasis and development of skeleton (Weitzel and Alexander Iwen, 2011, Singh et al., 2018, Zhang and Lazar, 2000). The biological effects of THs are cell- and tissue-specific, and are determined not only by a presence of hormones as such, but by many intra- and extracellular factors and circumstances, including transport across cell membrane, action of deiodinase enzymes, chromatin state, presence of thyroid hormone receptors and indispensable co-factors, and interplaying with other signals (Mancino et al., 2021, Zekri et al., 2022). The activity of HPT axis is environmentally sensitive, and THs act as a signal that ensures response of genome on the environmental changes, and thus shapes development in ecology-relevant context (Lema, 2020, Zwahlen et al., 2024). These circumstances allow considering the HPT axis as one of the fundamental evolutionary novelties of vertebrates, which ensured their rapid radiation (Norris (Norris and Carr, 2020, Esposito et al., 2021, Takagi et al., 2022), and put it among the foremost objectives of investigations not only in medicine, but also in developmental and evolutionary biology.

Teleosts, due to the high variety of skeletal tissues and the easy of manipulating with thyroid status, are extensively exploited models for the studies of the THs effects on vertebrate development. In the developing fish, THs content is not constant and displays pronounced rises and drops tightly linked in timing with the ontogenetic processes and life-history events, as well as circadian, ultradian, lunar and seasonal fluctuations (Grau, 1988, McMenamin and Parichy, 2013, Campinho, 2019, Deal and Volkoff, 2020, Kupprat et al., 2021). To study the role of THs in skeleton development, the model fishes with artificially altered hormonal status, THs deficiency or excess, are usually used. The TH deficiency is acquired via a transgene-mediated ablation of thyroid follicles and/or the pharmacological treatment of organism with the goitrogens - chemical agents suppressing synthesis of thyroid hormones or disrupting HPT axis. The effects of the excess of THs are being studied in transgenic and/or pharmacologically treated with THs animals (Shkil et al., 2012, Keer et al., 2019, Keer et al., 2022, Aman et al., 2023, Deal and Volkoff, 2020). Such studies have yielded a wealth of data of all sort of effects of THs on the development of cartilages, bones, scales and dentition in fish. Among others, the changes in the timing and rate of skeleton development can be identified as the major effects. THs deficiency commonly leads to postponed onset and retardation, while THs excess causes preliminary and accelerated development (Pelayo et al., 2012, Shkil et al., 2012, Keer et al., 2022). These developmental shifts might result in the pronounced phenotypic consequences, such as changes in the shape of skeleton elements and modules, number of serial structures, and even the loss of innate or appearance of atypical skeleton elements.

In the most of experimental models mentioned, the altered THs content is a persistent characteristic of the developing organism, and the observed phenotypic outcomes are essentially the cumulative consequences of multiple developmental shifts. However, many factors may have an acute effect on HPT axis function and cause transient fluctuations of THs content at any stage of development. What developmental and phenotypic outcomes these hormonal deviations can lead to, and which developmental stage(s) and skeleton structures are most sensitive to them, have been studied much less. Here, we present data on the developmental and phenotypic consequences of transient thyrotoxicosis and THs-deficiency at different stages of zebrafish, *Danio rerio* (Teleostei; Cyprinidae) skeleton patterning.

## Materials and Methods

### Fish and Experimental design

We conducted three experimental series to evaluate the effects of transient THs alterations at different developmental stages. For each series, we used a clutch of fertilized eggs obtained from the simultaneous spawning of 5-6 randomly selected pairs of “leopard” zebrafish, former *Brachydanio frankei* (5^th^-6^th^ generation from the fish collected in nature), provided by the aquarium department of the Severtsov’s Institute of Ecology and Evolution, RAS. To avoid fungal infections, eggs were treated with Methylene Blue. Then, each clutch was divided into seven groups: control (euthyroid, euTH) group and six experimental groups, the THs status of which was artificially changed at a certain developmental stage. The stages were determined in regards with the developmental milestones designated in the normal table of zebrafish postembryonic development (Parichy et al., 2009): stage 1, epiboly - hatching; stage 2, hatching - swim-bladder inflation; stage 3, swim-bladder inflation - appearance of caudal fin rays; stage 4, appearance of caudal fin rays - appearance of anal fin rays; stage 5, appearance of anal fin rays - appearance of pelvic fin buds; and stage 6, appearance of pelvic fin buds – formation of adult pigment pattern.

The groups were placed separately in the mesh cages with a stocking density ten individuals per liter (sd=10 ind/l), located in the main shared aquarium (V=300 l) with dechlorinated, UV-treated, aerated tap water, t=28^0^ C, 12/12 day/night regime, and daily replacement of water (1/5 the volume) (Supl. 1). To ensure the same temperature throughout the share aquarium, we used underwater water pumps. The newly hatched larvae were fed on TertraMin baby (Tetra, Germany) and NovoTom artemia (JBL, Germany). A week after the foraging onset, the diet was changed to TertraMin Junior (Tetra, Germany) and *Artemia salina* shrimps (Artemia Koral, Russia).

We provoked transient THs alterations in the traditional pharmacological way, administrating chemicals into the water at concentrations similar to those used effectively for zebrafish and other teleosts (Crane et al., 2006; Fetter et al., 2015; Cavallin et al., 2017; Shmidt et al., 2017; Roux et al., 2023). THs excess was induced by the treatment with 5 ng/ml of the exogenous triiodothyronine (T_3_), an active form of THs. THs deficiency was achieved by the administration of a mix of goitrogens (MPI *sensu* Salis et al. (Salis et al., 2021): Metimazol (300 µM/l) and potassium perchlorate (KClO_4,_ 7.2 µM/l) suppress T_4_ synthesis; and Iopanoic acid (1 µM/l) inhibits activity of iodothyronine deiodinases (DIO I-II), enzymes catalyzing T_4_-T_3_ conversion. For treatment, plastic aquarium (V=30 l) immersed in the share aquarium was used, so all conditions, with the exception of the chemical composition of the water, were similar among all groups. Totally, we conducted three experimental series (further series 1-3): in series 1 and 2, the experimental groups were treated with T_3_; and in series 3, the experimental groups were treated with MPI.

To determine the timing of onset and end of the treatment, we used key morphological and skeleton characters indicated by Parichy (Parichy et al., 2009)in the normal table of postembryonic zebrafish development. For this aim, we collected 5-10 live individuals, analyzed their general morphology, and then, stained them with green fluorescent marker of ossified tissues Calcein Am (Sigma) (Du et al., 2001) to analyze the skeleton patterning. If a change in THs status caused an acceleration or retardation of development, the experimental fish were treated until the appearance of at least several morphological characters indicating the end of the stage. After the treatment, we washed fish from the acting agents in a separate aquarium (V=100 l) with clean water for an hour. Then, fish were transferred back to the mesh cages in the shared aquarium, where they were reared until 100 days post fertilization (dpf).

### THs content analysis

In euTH fish, we assessed the developmental profiles of the THs: an active form – triiodothyronine (T_3_), and the prohormone – thyroxin (T_4_). For this aim, fish were collected daily from hatching to juvenile stage.

To study changes of content of the active form of THs (T_3_) in response to pharmacological treatment, we collected experimental fish in the middle of the each stage. To understand the dynamics of THs content after experimental treatment onset, we performed a 264 h experiment. For this aim, we used 7- and 32-dpf-old euTH fish (stages 3 and 6, respectively) and in each case divided them into three groups: control, T_3_-treated, and MPI-treated. The concentrations of chemical agents were the same as in the main experimental series. After 72 h, all groups were transferred to the clean water. During the 264 h experiment, fish were collected simultaneously in each group in the following mode: 0h/2h/4h/6h/8h/12h/24h/48h/72h/120h/168h/216h/264h.

To achieve a sample weight sufficient for analysis of the hormonal content (≥ 0.02 g), 50±5 individuals per sample were used at the early stages. Further, as the fish grew, the number of individuals per sample gradually decreased, but was at least ten. The intra-group variability of the THs content was out of scope of this study and each sample represented the average THs content of the group. The collected experimental fish were transferred to clean water for rinsing for 10-15 minutes, and then sacrificed with an overdose of 10% lidocaine anesthetic, washed in clean water, dried with filter paper and frozen in sample tubes (-24^0^ C). Using an enzyme-linked immunosorbent assay (ELISA), we assessed the content of total triiodothyronine (tT_3_) and total thyroxin (tT_4_) in the extract of whole fish. The extraction was performed in accordance with the Holzer et al. protocol (Holzer et al., 2017). Hormonal content was evaluated with the commercially available ELISA kits (Monobind Inc.) with a range 0.04–7.5 ng/mland 0.8-25 µg/dl for total T_3_ and T_4_, respectively. We used microstrip reader Stat Fax 303 plus (Awareness Technology) at 450 nm with reference wavelength at 620 nm in accordance with the manufacture’s protocol. The optical density was measured in duplicate for each sample (technical replication), and the mean value of the optical density was compared with standard curve to assess the actual hormonal concentration. Finally, hormonal concentration was recalculated to the sample weight and present in ng/g.

To test the interrelation of T_3_ and T_4_ developmental dynamics we used Spearmen’s rank correlation coefficient (r_s_). To compare the developmental dynamics of hormones between experimental series we implemented Kendall ranking coefficient method (Kendall τ). For better visualization of association between T_3_/T_4_ dynamics and similarity of the hormonal dynamics between different series, we used a polynomial trendline.

### Ossification pattern and morphological analysis

To assess the developmental effects of THs alterations, we collected euTH fish and experimental fish at the end of the stage. As were mentioned, 5-10 live individuals from each group were treated with green fluorescent marker Calcein Am. Additionally, 15-20 fish were sacrificed with overdose of 10% lidocaine, measured for standard length (SL) and fixed in 4% paraformaldehyde (PFA) for further staining for ossified tissues with Alizarine Red, bleaching and clearing in accordance with traditional methods (Walker and Kimmel, 2007). Terminology for bones follows Cubbage and Mabee (Cubbage and Mabee, 1996). For pairwise comparison of groups, we used parametric t-test.

To analyze the phenotypic consequences of treatment, we simultaneously euthanized all experimental fish (100 dpf), pictured them from the left side in a standard way, measured for standard length (SL, mm) and fixed in 4% PFA for further staining for ossified tissues with Alizarin Red. We estimated the number of: i) vertebrae; ii) soft rays in the paired and unpaired fins (upper and lower lobes in caudal fin were analyzed separately, as well as principal and procurrent rays); iii) branchiostegals; iv) infraorbitals; v) pharyngeal teeth; vi) supraneurals; and vi) number of scales in the lateral line row. We estimated minimal and maximal values (lim), mean (M), and standard deviations (SD). For pairwise comparison of groups, we used ANOVA and Tukey HSD Test. To visualize the variation of meristic traits and understand the main direction of changes, the groups were analyzed with Principal component analysis (PCA) for set of variables (number of vertebrae, rays in paired and unpaired fins, infraorbitals, branchostegals, supraneurals, pharyngeal teeth and scales).

To understand whether and how transient alterations of THs status affect developmental stability, we used fluctuating asymmetry (FA) as an indicator of developmental noise (Klingenberg, 2019, Zakharov et al., 2020). We assessed the FA of bilateral meristic traits most commonly used to study developmental stability in natural and laboratory populations of fishes: infraorbital bones count, number of soft rays in the paired fins, pharyngeal dentition formula, and number of scales in the lateral line row. The analysis was carried out on 100 dpf - old fish belonging to all experimental groups in each series. As a measure of FA, we used the average frequency of asymmetric manifestation per trait (Zakharov et al., 2020). For pairwise comparison of FA level between experimental groups, parametric t-test was implemented.

The statistics was performed with Statistica 6.0. We also analyzed the presence/absence and, if present, normal/abnormal morphology of the kinethmoid, proximal radials in the pectoral fin, hypurals and epural in the caudal fin, and structures composing the Weberian apparatus.

Landmark-based geometric morphometrics was implemented to compare the head shape of the 100 dpf fish. The position of eleven morphological landmarks (eight homologous landmarks and three semilandmarks defining a slope of the craniofacial profile) (Supl. 2) were digitize from pictures with TPSdig v2.0 (Rohlf, 2015). To clarify the shape differences, we implemented Canonical variate analysis (CVA), and Discriminant function after the Procrustes ANOVA, and a pairwise Mahalanobis distances were calculated with 10,000 permutation rounds (Klingenberg and Monteiro, 2005). For better visualization of shape differences, we created a wireframe graph. The analysis was performed with MorphoJ v1.06d (Klingenberg, 2011).

## Results

### Developmental dynamics of THs content in the control and experimental groups

In euTH fish, growth rate, as well as sequence and timing of developmental events were similar, and did not differ from those described for zebrafish by Parichy et al. (2009). The developmental dynamics of T_3_ and T_4_ content were highly correlated (r_s_=0.68-0.933, p [ 0.0001) in euTH fish, and did not differ between series (τ=0.507-0.515, p [ 0.001). Their THs profiles were characterized by an increase at the early developmental stages and a subsequent decline until the stage 6, at which a short-term rise of T_3_ content occurred in all series (Fig. 1).

**Fig. 1.**
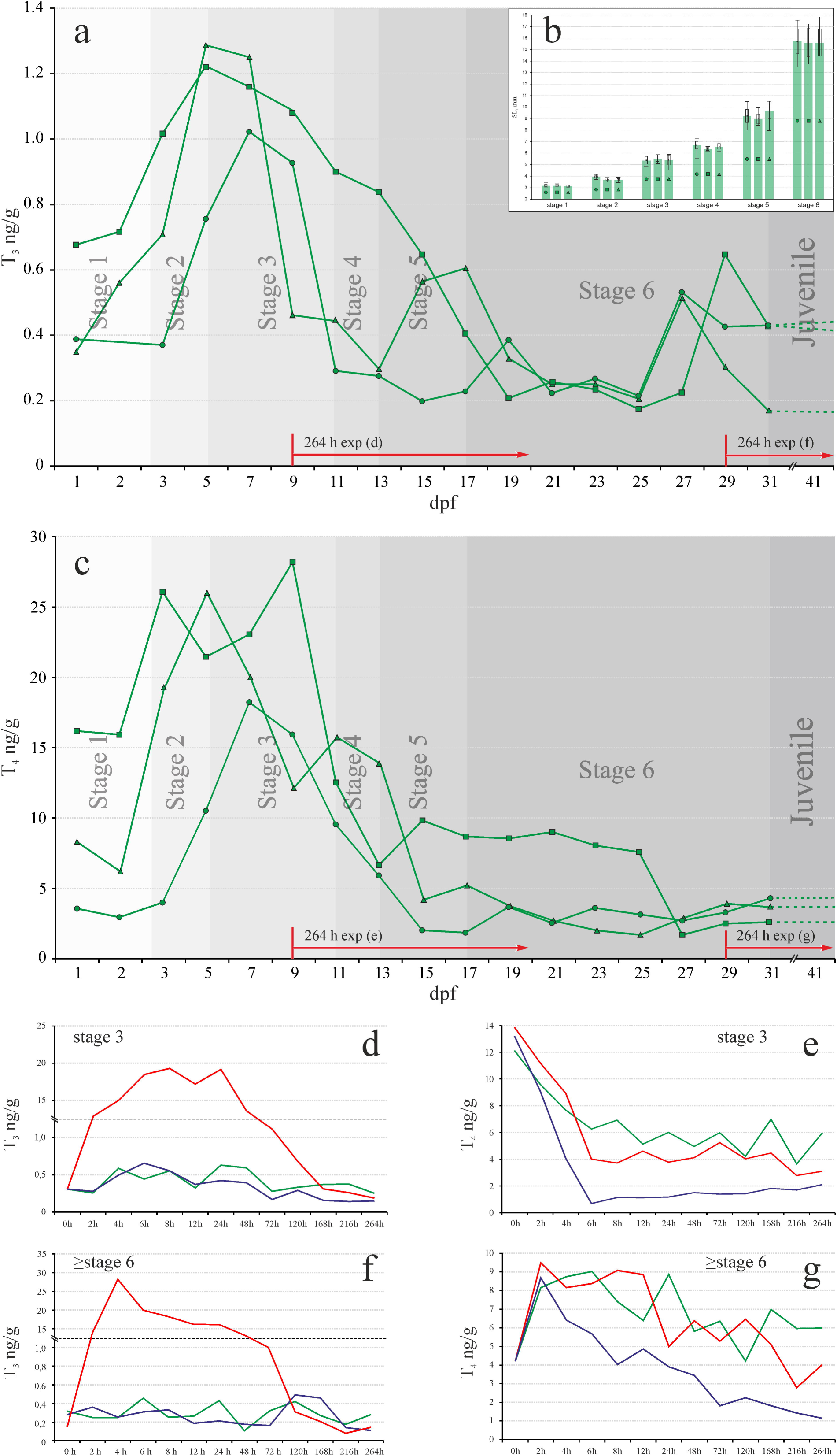
Developmental dynamics of thyroid hormones: (a/c) - T_3_/T_4_ ng/g, respectively, and (b) - standard length (SL, mm) in euthyroid zebrafish (series 1-3), and (d-g) in the 264 h experiments. Stage 1- 6 developmental stages. dpf – days post fertilization. 264 h exp – timing of 264 h experiment.

In the middle of each stages, the high values of T_3_ content were detected in all hormonally treated group. In contrast, MPI-treated groups did not display significant changes in T_3_ level (Supl. 3). The 264 h experiments also revealed a stable T_3_ content in euTH and MPI fish (Fig. 2). Moreover, the 264 h experiment demonstrated that at different developmental stages T_3_ treatment did not affect T_4_, but led to a rapid increase of T_3_ content in fish, which persisted for up to 48 hours, and then, after the transfer to the clean water, returned relatively quickly to euTH values. This finding indicated that hormonal treatment caused a transient T_3_ excess or thyrotoxicosis *sensu* De Leo et al. (De Leo et al., 2016). In contrast, MPI treatment did not change T_3_, but somewhat decreased and mitigated fluctuations of T_4_ level (Fig. 1). This effect appeared in the first 6-8 hours after the treatment onset and persisted throughout experiment, i.e. led to the T_4_ deficiency.

**Fig. 2.**
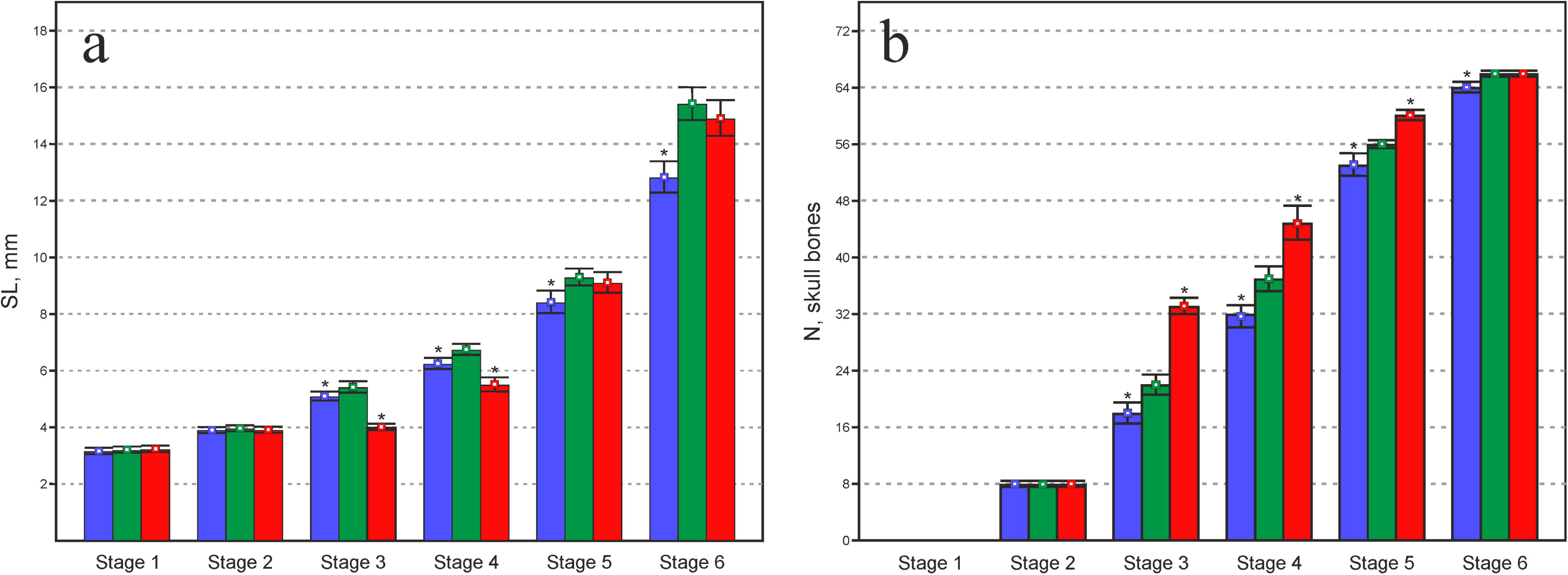
Standard length (SL, mm) and number of skull bones (N, skull bones) in experimental groups: control (green); T_3_-treated (red); and MPI-treated (blue). Asterisks indicate statistically significant differences from the control group. Whiskers indicate the standard deviations (SD).

### Developmental effects of THs alterations

#### Stage 1. Epiboly – hatching

We failed to find differences in the developmental timing, size and skeleton pattern among the experimental groups (Fig. 2). All newly hatched larvae possessed the cleithrum, a membranous bone in the pectoral fin girdle (Fig. 3a).

**Fig. 3.**
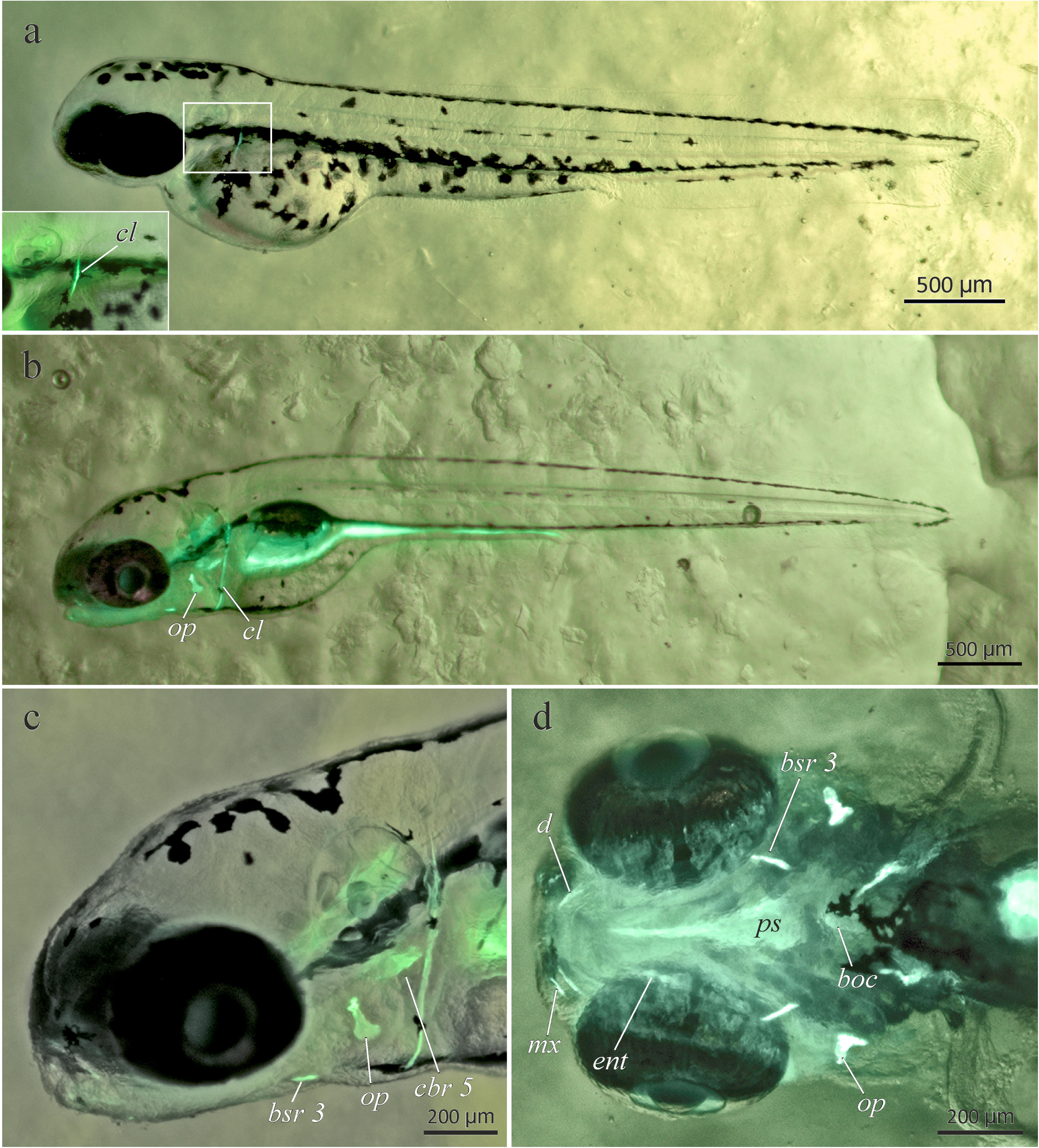
Ossified structures in the control, T_3_- and MPI-treated fish at the end of stage 1 (a) and stage 2 (b-d). a, b – total view, c – lateral view of head, d – ventral view. boc – basioccipital, bst 3 – branchiostegal 3, cbr 5 – ceratobranchial 5, cl – cleithrum, d – dentary, mx – maxilla, ps – parashenoid.

**Fig. 4.**
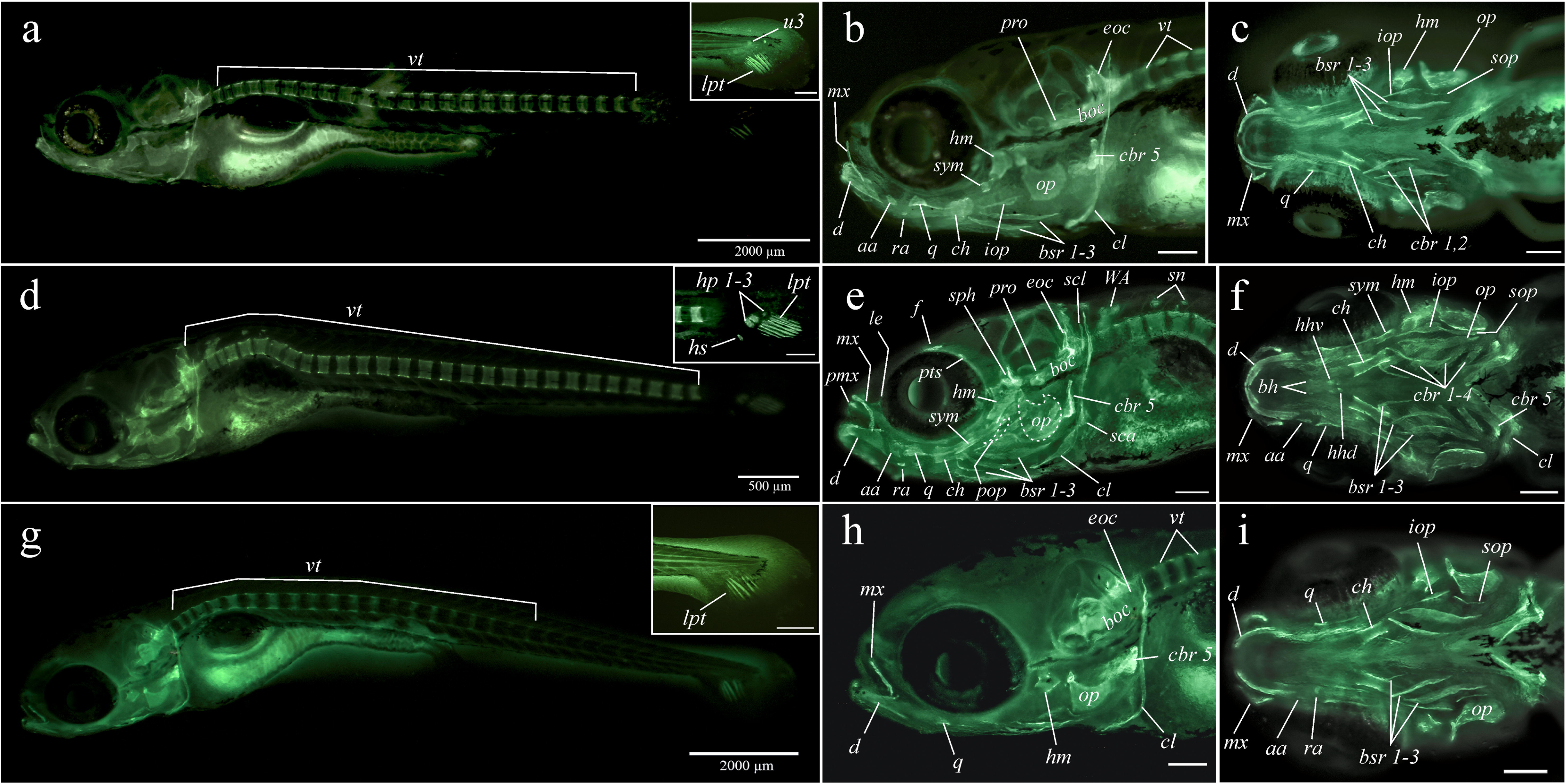
Ossified structures in the control (a-c), T_3_- (d-f) and MPI-treated (g-i) fish at the end of stage 3. a, d, g – total view; b, e, h – skull lateral projection; c,f,i – skull ventral projection. Cranial ossifications: aa – anguloarticular; bh – basihyal; boc – basioccipitale; bst 1-3 – branchiostegals 1-3; cbr 1-5 – ceratobranchials 1-5; ch – ceratohyal; d – dentary; ent - entopterygoid; eoc – exooccipital; f – frontal; hhd/hhv – hypohyal dorsal/ventral; hm - hyomandibula; iop – interopercula; leth – lateral ethmoid; mx – maxilla; op – opercula; pro - prootic; ps – parashenoid; pmx - premaxilla; pop – preopercula; pts – pteroshenoid; q- quadrate; sop – subopercula; sph – sphenotic; sym - symplectic. Postcranial ossifications: cl – cleithrum; hyp 1-3 hypurals 1-3; lpt – lepidotrichia (fin rays); sca – scapula; scl – supracleithrum; sn – supraneurals; u3 – ural 3; vt – vertebrae; WA – Weberian apparatus.

#### Stage 2. Hatching – gas bladder inflation

In the experimental groups, differences in developmental timing, size and skeleton pattern were not observed (Fig. 2). The vast majority of larvae developed the skull bones: parasphenoid, basioccipital, dentary, maxilla, ceratobranchial 5, branchiostegal 3, opercula and entopterygoid (Fig. 3b-d). A few larvae, regardless of treatment, also possessed ceratohyal, hyomandibula, and quadrate.

#### Stage 3. Gas-bladder inflation - appearance of rays in caudal fin

The duration of the stage did not differ between euTH and T_3_-fish (≈120-122 h), but was extended to 200 h in the MPI-group, that along with the significant deceleration in growth indirectly indicated the effects of THs deficiency on the fish. In T_3_-fish, growth was arrested. As a result, at the end of the stage, euTH and experimental larvae significantly differed in size (p<0,001) (Fig. 2a).

They also displayed discrepancies in the skeleton pattern (Fig. 2b; 4). At the end of the stage, most of euTH fish possessed 22 skull ossifications: four neurocranial (parasphenoid, basioccipital, exoccipital, and prootic), and 18 splanchnocranial bones (dentary, maxilla, anguloarticular, retroarticular, quadrate, hyomandibula, symplectic, ceratohyal, ceratobranchial 1,2,5, entopterygoid, branchiostegal rays 1-3, opercula, subopercula, and interopercula). T_3_-fish demonstrated an advanced state of the skull development and possessed 33 skull ossifications: eight neurocranial (four euTH bones plus lateral ethmoid, frontal, pterosphenoid and sphenotic bones), and 25 splanchnocranial (18 euTH bones plus praemaxilla, basihyal, hypohyal ventral and dorsal, ceratobranchial 3 and 4, and praeopercula). The bones similar to those in the euTH fish were at advanced state. Compared to the euTH fish, MPI-fish demonstrated retarded development and had only 18 less ossified skull bones. They did not develop the prootic bone in the neurocranium; as well as ceratobranchials 1-2 and symplectic in the splanchnocranium.

The postcranial skeleton differed too (Fig. 2b; 4). The euTH fish had 24-25 vertebrae, cleithrum in the pectoral girdle, weakly ossified ural 3+4 centrum and three-four principal rays in each lobes of the caudal fin. T_3_-fish had 27-28 vertebrae, and many of them displayed a curvature (kyphosis) at the anterior portion of precaudal (abdominal/thoracic) part of the vertebral column. They also possessed abnormal scaphium and supraneural 2 in the Weberian apparatus, as well as ossified structures located dorsally to the fourth and fifth vertebrae (presumably supraneurals 4 and 5). Several T_3_-fish developed the tripus and rib 4. The pectoral girdle of T_3_-fish consisted of the cleithrum, supracleithrum and scapula. The hypurals 1-3 and ten principal fin rays, five in each lobes were present in the caudal fin. MPI-treated fish had 19-20 vertebrae, cleithrum in the pectoral girdle and 5-6 rays in the caudal fin (3 and 2 or 3 and 3 in the lower and upper lobe, respectively).

#### Stage 4. Caudal fin rays - anal fin rays

The duration of the stage (≈100 h) was the same in the euTH and T_3_-groups. In the MPI-group, the stage lasted for 130 h. The euTH fish demonstrated a steady growth; MPI-group displayed a moderate slowdown; and in T_3_- group growth rate was significantly decreased. As a result, at the end of the stage, larvae significantly (p<0,001) differed in size (Fig. 2a).

The euTH fish displayed an intensive development of the ossified skull. At the end of the stage, the majority of euTH fish had 37 bones (Fig. 2b; 5). The lateral ethmoid, frontal, sphenotic, pterosphenoid, pterotic and posttemporal appeared in the neurocranium. The praemaxilla, coronomeckelian, epihyal, urohyal, hypohyal dorsal and ventral, basihyal, ceratobranchials 3-4 emerged in the splanchnocranium. Compared to euTH fish, T_3_-fish demonstrated an advanced state of the skull development and possessed 45 skull bones. They differed from the euTH fish in the presence of supraoccipital in the neurocranium, as well as epibranchials 1-4, ecto- and metapterygoid, and lacrimal in the splanchnocranium. Most of the bones seemed more ossified, massive and robust, and had another shape than those in euTH fish. MPI-fish had only 32 less ossified than in euTH fish skull bones. They did not have the pterosphenoid, posttemporal, coronomeckelian, epihyal and lateral ethmoid.

In all experimental groups, all vertebrae were ossified and possessed the neural and haemal spines (Fig. 5). All groups had the ossified Weberian apparatus. T_3_-fish also developed the partial ossification of supraneurals 2-3. The number of ribs differed among the groups: euTH fish had 4-5 ribs; T_3_-treated – 6-7 ribs; and MPI-treated – 2-3 ribs. The caudal fin complex in all groups possessed preurals 2-3, pleurostyle, hypurals 1-5, parhypural, and 8-9 principal rays in each lobe. The euTH and and a few MPI-individuals, had a single procurrent fin ray in the lower lobe. T_3_-fish developed a procurrent ray in both lobes. The euTH and MPI-fish possessed the fused urals 1-2, and ossified in various degree ural 3. In T_3_-fish, ural 3 was not found. In the dorsal fin, euTH and T_3_-fish developed 2-3 rays. In the anal fin, all larvae had 6-7 soft rays. The composition of the pectoral fin and girdle differed among groups. Presence of cleithrum, supracleithrum and initial ossification of scapula cartilage was characteristics of euTH fish. T_3_- fish were more advanced and had the postcleithrum, scapula, coracoideum and 4-5 ossified rays. The absence of scapular ossification was typical of the MPI-fish.

**Fig. 5.**
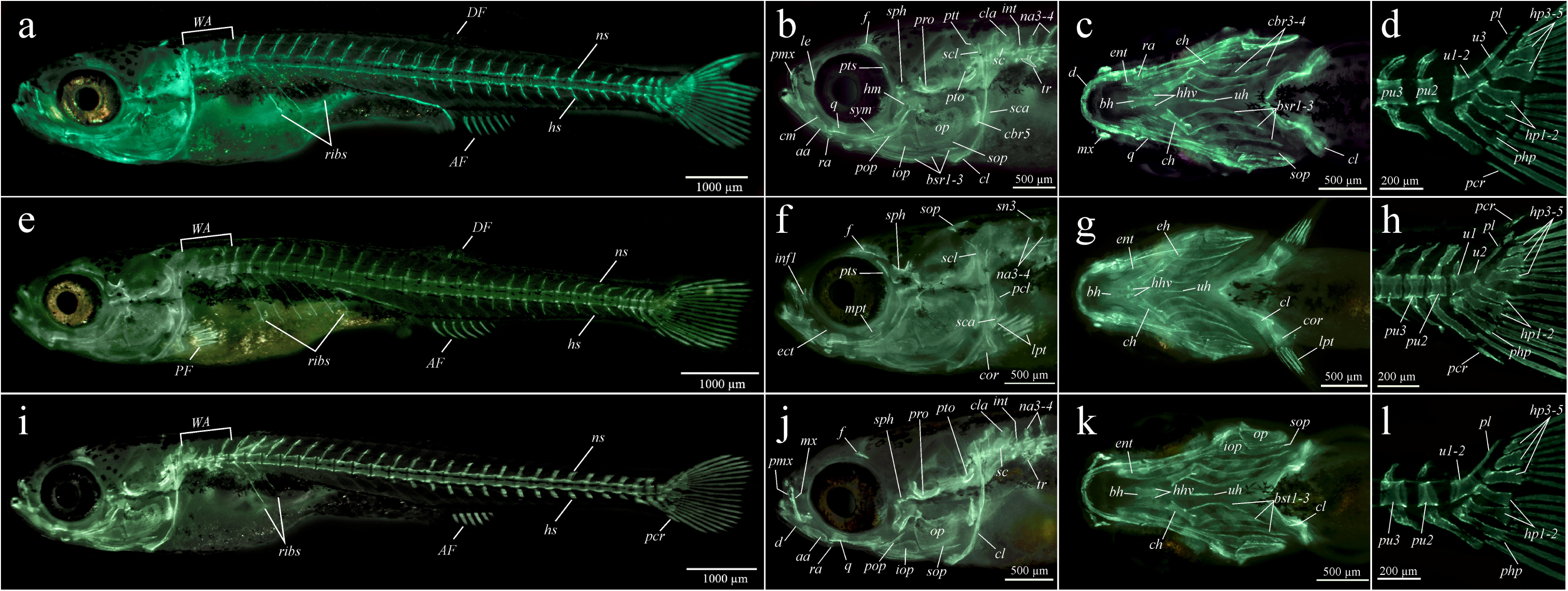
Ossified structures in the control (a-c), T_3_ – (d-f) and MPI-treated (g-i) fish at the end of stage 4. a, e, i – total view; b, f, j – skull lateral projection; c, g, k – skull ventral projection; d,h,l – caudal fin. Cranial ossifications: aa – anguloarticular; bh – basihyal; bst 1-3 – branchiostegals 1-3; cbr 5 – ceratobranchial 5; ch – ceratohyal; cm – coronomeckelian; d – dentary; ect – ectopterygoid; ent - entopterygoid; eh - epihyal; f – frontal; hh – hypohyals; hm - hyomandibula; iop – interopercula; leth – lateral ethmoid; mpt – metapterygoid; mx – maxilla; op – opercula; pro - prootic; pto – pterotic; pmx - premaxilla; pop – preopercula; pts – pteroshenoid; ptt – postemporal; q- quadrate; sop – subopercula; sph – sphenotic; sym – symplectic; uh - urohyal. Postcranial ossifications: AF/DF/PF – anal, dorsal and pectoral fins, respectively; cl – cleithrum; cla – claustrum; cor – coracoid;hs – haemal spine; hyp 1-5 hypurals 1-5; int – intercalarium; lpt – lepidotrichia (fin rays); na3-4 – neural arches 3-4; ns – neural spine; pcl – postcleithrum; pcr - procurrent fin ray; php – parhypural; pl – pleurostyle; pu2-3 preurals 2-3; ribs – ribs; sc – scaphium; sca – scapula; scl – supracleithrum; sn3 – supraneural 3; tr – tripus; u1-3 – urals 1-3; WA – Weberian apparatus.

#### Stage 5. Anal fin rays - pelvic fin buds

The duration of the stage was the same in euTH and MPI-fish (≈90-96 h), but was reduced in T_3_- fish (≈48 h). This period was characterized by an active growth of euTH and T_3_-fish. Growth of MPI-fish was retarded that resulted in significant difference in size (p<0,001) with the euTH-fish at the end of the stage (Fig. 2a).

In euTH fish, the ossified skull demonstrated a notable progress. At the end of the stage, the majority of them had 56 skull bones (Fig. 2b; 6). The supraoccipital, supraorbital, vomer, orbitosphenoid, epiotic, parietal, supra- and kinethmoid appeared in the neurocranium. The meta- and ectopterygoid, epibranchials 1-4, basibranchials 1-3, pharyngobranchial 2+3, and palatine emerged in the splanchnocranium. In T_3_-fish, a less pronounced than at previous stages, but significant acceleration of the skull development occurred. At the end of the stage, they had 60- 61 skull bones, including the “late” bones such as infraorbital 3 (a few individuals had even infraorbital 4), pharyngobranchial 1, mesethmoid and nasal. Generally, the head of T_3_-fish was relatively shorter and more “heavy” than in euTH-group. In contrast, MPI-fish displayed a retardation of the skull development, and usually possessed 53 bones only. They did not develop kinethmoid, supraethmoid and supraorbital. The bones present were less ossified than those in euTH fish.

All experimental groups had a complete pattern of ossifications in the vertebral column and associated with vertebrae elements (ribs, neural and haemal arches and spines); full pattern of ossicles in the Weberian apparatus and unpaired fins; ossified supraneurals and intermuscular bones (Fig. 6). They did not display discrepancies in the pelvic fin and girdle, but differed in the state of the pectoral fin and girdle. Thus, in euTH fish, girdle was composed by the cleithrum, supracleithrum, postcleitrum, and partially ossified scapula and coracoideum. The fin possessed two ossified proximal radials and 10-12 rays. T_3_-fish differed from euTH group by the presence of mesocoracoideum, advanced ossification of the scapula and coracoideum, and abnormal proximal radials. MPI-fish displayed weakly ossified scapula and coracoideum, absence of radials, and a fewer number of rays in comparison with euTH fish.

**Fig. 6.**
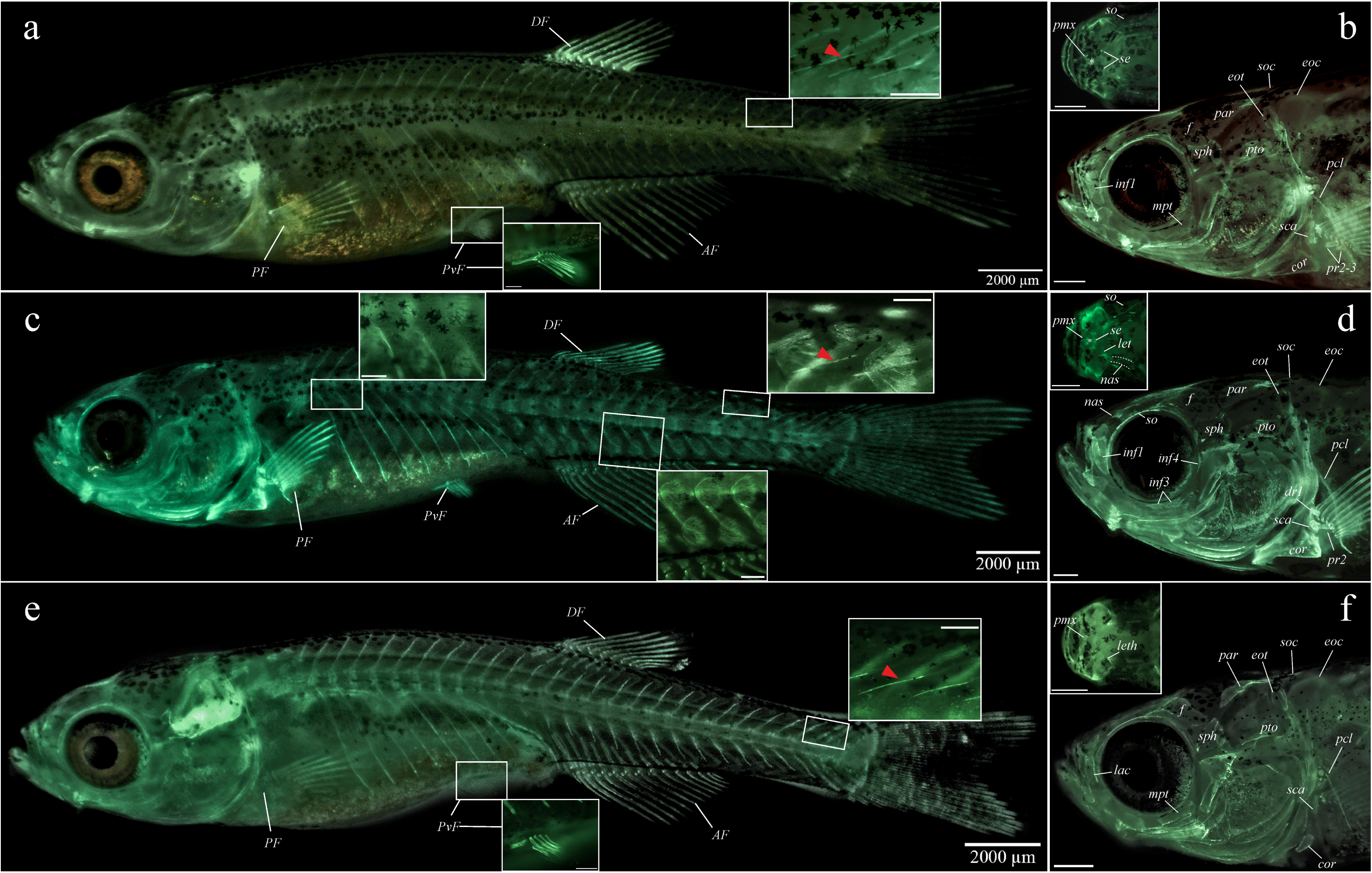
Ossified structures in the control (a-b), T_3_- (c-d) and MPI-treated (e-f) fish at the end of stage 5. a,c,e – total view; b,d,f – skull lateral projection. Cranial ossifications: eoc – exooccpital; eot – epiotic; f – frontal; hh – hypohyals; hm - hyomandibula; inf3-4 – infraorbitals 3-4; iop – interopercula; lac – lacrimal (infraorbital 1); leth – lateral ethmoid; mpt – metapterygoid; na – nasal; par – parietal; pto – pterotic; pmx - premaxilla; seth – supraethmoid; so – supraorbital; soc – supraoccipital; sph – sphenotic; * - kinethmoid. Postcranial ossifications: AF/DF/PF/PvF – anal, dorsal, pectoral and pelvic fins, respectively; cor – coracoid; dr – distal radial; pr 2-3 – proximal radials; pcl – postcleithrum; sca – scapula; red arrow – intermuscular bones.

In several euTH fish, early development of squamation (one-two rows composed by several, less than ten, weakly ossified scales at the caudal peduncle) was detected. Fish treated with T_3_ had squamation covering almost the entire body. MPI-fish were scale-less (Fig. 6).

#### Stage 6. Pelvic fin buds - adult pigment pattern

The duration of this stage was similar for all experimental group. The euTH and T_3_-fish did not differ in growth rate. MPI-fish displayed a growth retardation, and at the end of the stage, they were significantly smaller (p<0,001) than euTH individuals (Fig. 2a).

The euTH and T_3_-treated group developed almost all bones of the adult skeleton, including infraorbital 4 and 5, anterior and posterior sclerotic, hypobranchial 3 and squamation (Fig. 7). Only the most «developmentally late» bones: hypobranchial 1 and 2, intercalar, preethmoideum and infraorbital 2 were absent. A few T_3_- fish (≈10% of individuals) had infraorbital 2. The skeleton pattern of MPI-fish was truncated in comparison with euTH fish. The vast majority of them did not developed infraorbital 4 and 5, and about a quarter of them (≈25% of individuals) - infraorbital 3. The squamation development was also incomplete; the abdomen was not covered by scales (Fig. 7).

**Fig. 7.**
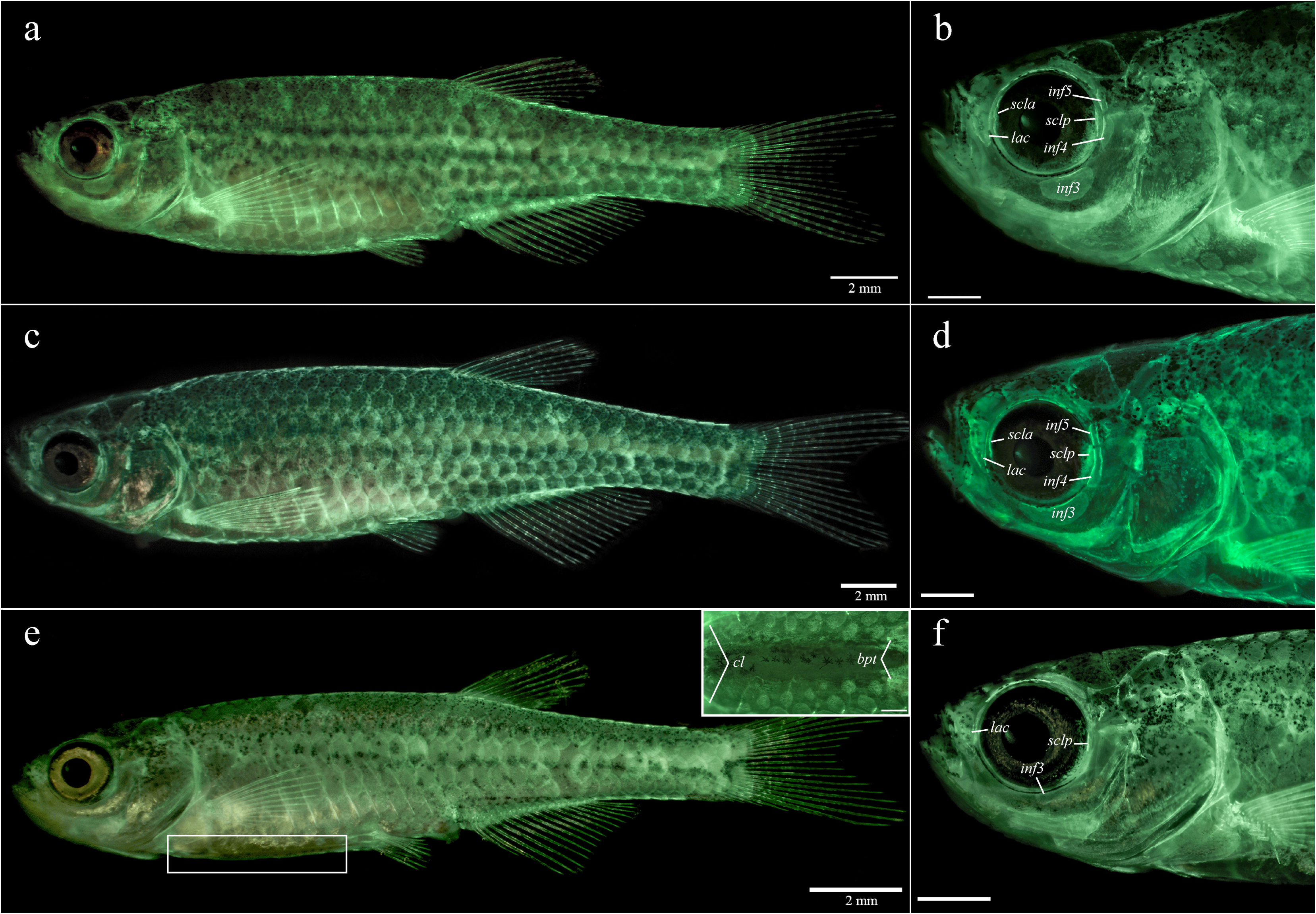
Ossified structures in the control (a-b), T_3_- (c-d) and MPI-treated (e-f) fish at the end of stage 6. a,c,e – total view; b,d,f – skull lateral projection. Cranial ossifications: inf 3-5 – infraorbitals 3-5; lac – lacrimal (infraorbital 1); scla/sclp – sclerotic anterior/posterior. Postcranial ossifications: cl – cleithrum; bpt – basipterigium

### The phenotypic consequences of T_3_ and MPI-treatment at the different developmental stages in 100 dpf old fish

#### Adult morphology in control groups

Despite the fact that variability of meristic traits and head morphology of euTH fish (100 dpf old) did not go beyond the species-specific reaction norm, in the entirety of meristic characteristics, as well as in head shape, fish belonging to different series differed (Supl. 4-5). The most pronounced discrepancies were found between series 1 and 3, whereas in head shape – between series 2 and 3 (Mahalonobis distance = 11.028, p[0,001; Procrustes ANOVA F=9.84 p [0,0001, SS ind/res=0.0280/0.0556). Taking into account that intraspecific differences in meristics and morphology might have various background, including genetics (Ferreri et al., 2000, Shkil et al., 2018, Martini et al., 2021), the analysis of the phenotypic consequences of experimentally induced THs alterations was conducted for each line separately.

##### Stage 1. Epiboly – hatching

We failed to find noticeable developmental shifts in response to T_3_ and MPI-treatment at the stage 1. However, T_3_-treated fish demonstrated a small but statistically significant (p≤0.03) decrease in the number of lateral line scales in comparison with the euTH group. Moreover, in series 1, T_3_-fish also displayed a modest decrease in the count of pharyngeal teeth (p≤0.02), and in series 2 - pectoral fin rays (p≤0.02) (Fig. 8). The differences in head morphology and FA level were not revealed between euTH and T_3_-fish, but found between euTH and MPI-fish (Fig. 9).

**Fig. 8.**
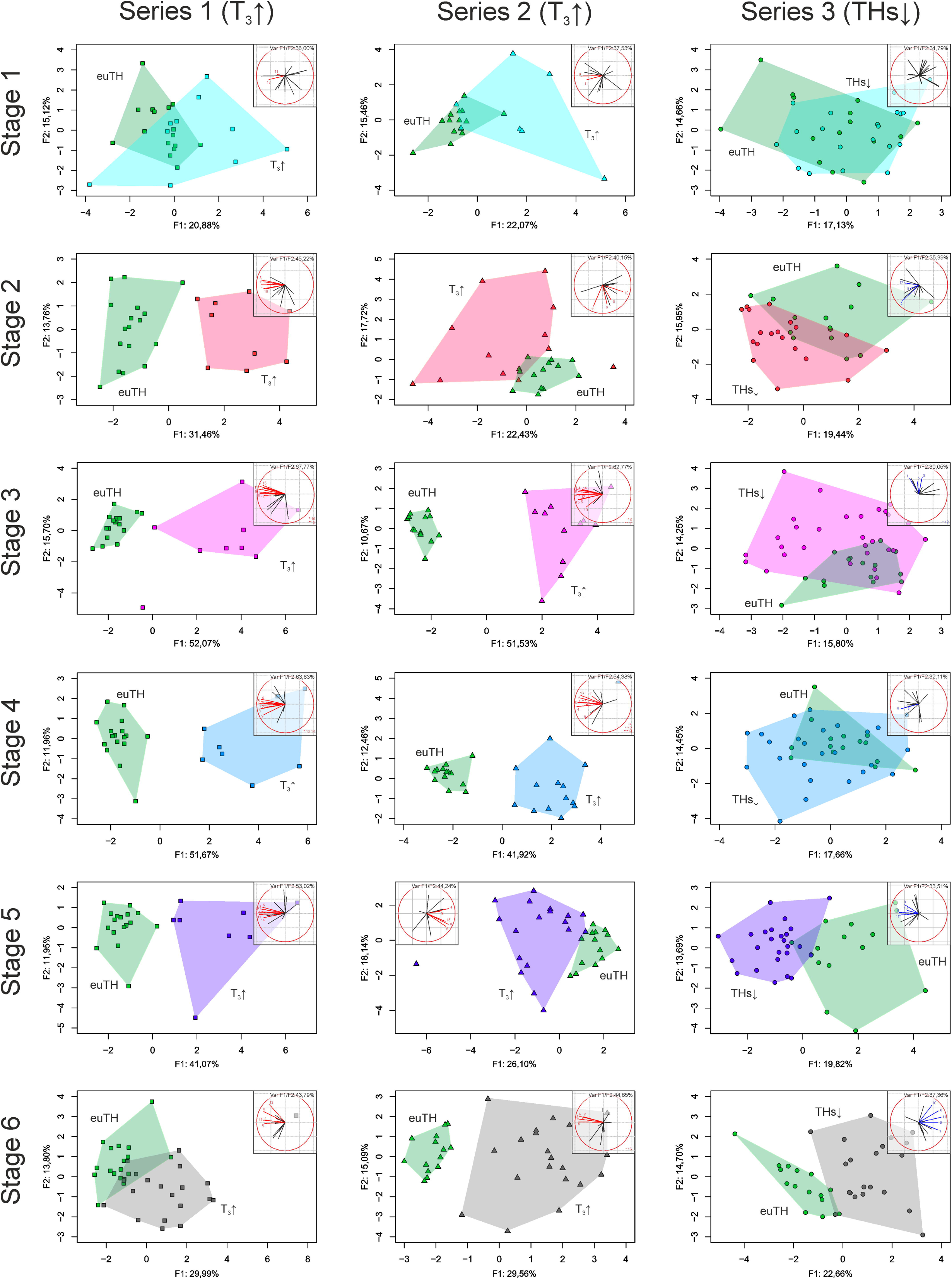
Phenotypic consequences of hormonal and goitrogen treatment at the stage 1-6 in series 1-3. Principal component analysis of the control (euTH) and experimental groups (T_3_↑ - thyrotoxicosis, and THs↓ - THs-deficiency) based on the entirety of meristic traits with grouping of the variables in two main principal components (Correlation circle). Characters displayed significant differences are marked by colors (red – in T_3_ treated groups, blue – in MPI treated group). 1 – vertebrae; 2 – dorsal fin rays; 3 – anal fin rays; 4 – caudal fin procurrent rays (upper); 5 – caudal fin principal rays (upper); 6 – caudal fin procurrent rays (lower); 7 – caudal fin principal rays (lower); 8 – pectoral fin rays; 9 – pelvic fin rays; 10 – infraorbitals; 11 – scales in lateral line row; 12 – branchiostegals; 13 – pharyngeal dentition; 14 – supraneurals.

**Fig. 9.**
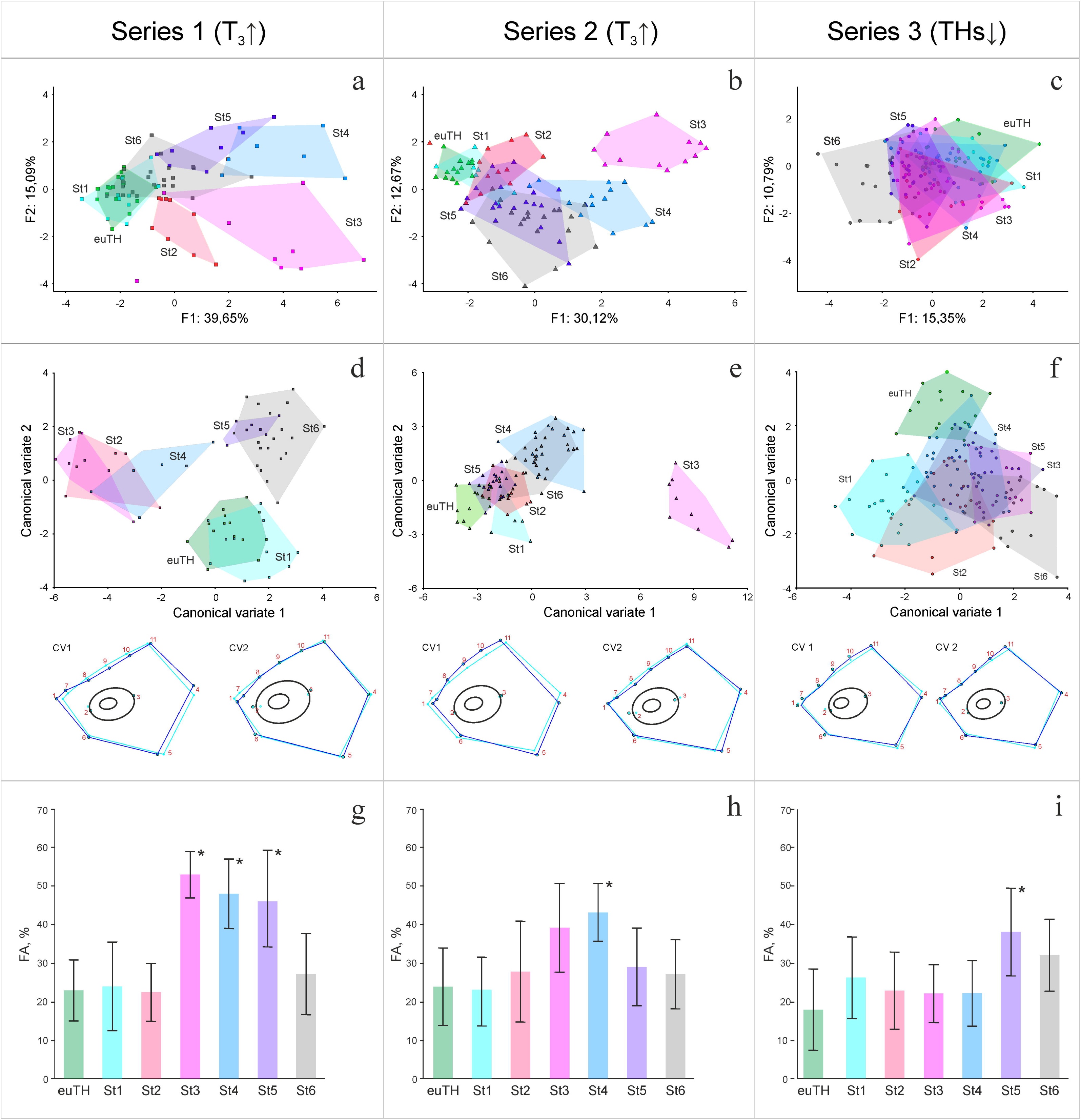
Phenotypic consequences of hormonal and goitrogen treatment in the experimental series 1-3. a-c - Principal component analysis of meristic traits, d-f - Canonical variate analysis (CVA) of the head morphology, and g-i -fluctuating asymmetry level (FA - frequency of FA in meristic traits, %). St1-6 – experimental groups treated at the stages 1-6. * indicates statistically significant differences in FA level between experimental groups and euTH fish.

##### Stage 2. Hatching – swim-bladder inflation

Despite an absence of the pronounced developmental reaction on the hormonal and MPI- treatment at this stage, the phenotype of experimental fish differed from the phenotype of euTH group (Fig. 8-9). The T_3_-fish possessed the decreased number of infraorbitals, lateral line scales, and pectoral fin rays. They also displayed changes in the Weberian apparatus. The majority of fish displayed the shortened lateral processes of vertebrae 1, an absence of intercalarium, the abnormal scaphium and rib 4, and reduction in size of the supraneurals 2 and 3. Approximately a half of T_3_-fish had an abnormal reduced in size kinethmoideum. Several fish had the reduced in number or/and abnormal proximal radials of the pectoral fin. We revealed differences in the reaction on the T_3_ between the series 1 and 2 (Fig. 8). Series 1 displayed a reduction in the number of pharyngeal teeth and vertebrae, whereas series 2 did not, but had fewer procurrent caudal fin rays. The geometric morphometric revealed changes in the head shape in series 1 only, fish were characterized by a sloping head profile and shortened preorbital portion of the skull in comparison with euTH.

MPI-fish had the significantly increased count of soft rays in the pectoral and dorsal fins, as well as the increased number of vertebrae and branchiostegals. They statistically differed in the head morphology but did not in the FA level from the euTh group (Fig. 9).

##### Stage 3. Swim-bladder inflation – appearance of rays in caudal fin

The hormonal treatment led to numerous phenotypic consequences (Fig. 8-9). Only the number of unpaired fin rays and branchiostegal rays in series 1, and the number of vertebrae, dorsal and anal fin rays, and principal rays of the caudal fin in series 2 did not differ from euTH fish. The count of the remaining meristic characters was significantly reduced. Moreover, T_3_-fish demonstrated numerous reductions and malformations in the Weberian apparatus. In particular, they displayed an absence of intercalarium and lateral processes of vertebra 1. The majority of fish did not develop claustrum, and had the malformed scaphium, tripus, supraneural 2, supradorsals 3 and 4, and rib 4. Many of T_3_-fish demonstrated an absence or reduction of the kinethmoid, reduction in the number and malformations of proximal radials in the pectoral fin, complete or partial absence of pelvic fin and girdle, loss of hypural 6 and epural in the caudal fin, as well as an abnormal shape of parhypural.

The overall morphology of the T_3_-treated fish drastically differed from euTH group. They were significantly smaller (p[0.001), and many of them had curvature(s) of the vertebral column, the deformed jaws and preorbital portion of the neurocranium. Several fish did not display pronounced skull malformations, but in spite of this, significantly differed in the head shape from the control group. They displayed a sloping head profile due to abnormally developed preorbital portion of skull. In both series, FA level in T_3_-fish was significantly higher than in the euTH group (Fig. 9).

The phenotypic consequences of MPI-treatment were much weaker. MPI-fish had the increased number of vertebrae, branchiostegals, and soft rays in pectoral and dorsal fins. They also differed from euTH fish in the shape of a head, but the differences seem biologically negligible (Fig. 9).

##### Stage 4. Rays in caudal fin – anal fin rays

The hormonal treatment led to numerous outcomes in adult phenotype (Fig. 8-9). The number of the most of meristic characters was reduced. In both series, the infraorbitals, anal fin rays and principal rays in lower lobe of the caudal fin were not affected. We found differences in reactions between series. Thus, in series 1, the principal rays in the upper lobe of the caudal fin and branchiostals; and in series 2 - vertebrae and dorsal fin rays did not change in comparison with euTH group. The hormonal treatment also led to deformities in the Weberian apparatus elements: lateral proc. of vertebra 1, supraneural 2, supradorsal 3 and 4. Several fish did not develop the intercalarium. We revealed the numerous malformations of the precaudal and caudal vertebrae and associated elements (ribs, neural and haemal arches). In the pectoral fin, the number of proximal radials was reduced, and the radials were abnormal. Several fish did not possess pelvic fin and girdle at one or both sides. A few fish displayed an absence of epural and deformed parhypural in the caudal fin. The third part (≈30%) of individuals did not develop the kinethmoid bone. The remaining fish had a reduced kinethmoid. FA level was significantly higher than in euTH groups. The geometric morphometrics revealed significant differences between T_3_ and euTH fish. The former had a shortened head due to a reduction of the preorbital portion of the skull. MPI caused a significant increase in the number of rays in the pectoral and pelvic fins. FA level and shape of the head were similar to euTH fish (Fig. 9).

##### Stage 5. Anal fin rays – pelvic fin bud

Treated with T_3_ at this developmental stage fish displayed a significant reduction in the number of pharyngeal teeth, procurrent and soft rays in the caudal and pectoral fins, respectively (Fig. 8). Series 1 also demonstrated a decrease in the count of lateral line scales and supraneurals. Series 2 possessed the reduced number of pelvic fin rays. Approximately a third of individuals belonging to both series was characterized by the reduction in size of the lateral processes of vertebra 1 and dorsal process of neural complex in the Weberian apparatus. About a half of fish displayed malformations in the pectoral fin: reduction in the number and various deformities of proximal radials. In series 1, FA level of T_3_-fish was significantly higher, whereas in series 2 – comparable with euTH fish. The same tendency was revealed for the head shape: T_3_-fish belonging to the series 1 differed from euTH fish, whereas T_3_ fish belonging to the series 2 did not (Fig. 9).

MPI treatment resulted in a significant increase in the number of infraorbitals and lateral line scales, as well as soft rays in the pectoral and pelvic fins. FA in MPI-fish was significantly higher than in euTH fish. The shape and proportion of head did not differ from the reference group (Fig. 9).

##### Stage 6. Pelvic fin buds – adult pigment pattern

The vast majority of adult hormonally treated fish possessed the significantly reduced number of pharyngeal teeth, and fin rays: procurrent - in the caudal; and soft – in pectoral fin (Fig. 8). They also were characterized by the loss and/or malformation of proximal radials in the pectoral fin. Moreover, series 1 demonstrated a decrease in the number of lateral line scales. Series 2 had the reduced count of pelvic fin rays. FA level in fish belonging to both series was comparable with euTH fish. Head morphology of T_3_- fish differed from euTH due to slight change in a slope of the skull profile (Fig. 9).

MPI-fish had the increased number of infraorbitals, lateral line scales, and fin rays: principal – in caudal, soft - in pectoral and pelvic. FA level was significantly higher than in euTH fish. The head shape of MPI-fish differed significantly from euTH group (Fig. 9).

## Discussion

THs developmental profiles in euTH fish displayed two surges: the first and most pronounced occurred before the transition stage, and the second preceded the formation of the adult pigment pattern. Treatment with T_3_ and MPI caused transient acute thyrotoxicosis and THs deficiency, which affected the skeleton development: led to the acceleration and developmental delay, respectively. Bones differed in THs-sensitivity. In response to changes of THs status, some of them displayed a notable shift in the developmental timing and rate, whereas other demonstrated a subtle or absence of reaction. The developmental stages differed in THs sensitivity too. The embryonic and early larval stages should be regarded as the least sensitive; two subsequent stages - the most sensitive; and further development demonstrated a moderate sensitivity. Transient thyrotoxicosis and THs deficiency resulted in various phenotypic consequences: changes in the number of meristic characters, the shape and composition of skeleton complexes and individual structures.

### THs dynamics in euTH and experimental fish

Despite the extensive use of zebrafish as a model species for THs research (Marelli and Persani, 2017, Lazcano et al., 2023), studies of THs developmental dynamics remain fragmented and focus on the certain, mainly early, developmental stages. However, due to them, we know that zebrafish eggs contain maternal THs, which is consumed during embryonic development, and the functional HPT axis is formed shortly after the hatching (Porazzi et al., 2009, Walpita et al., 2013, Chang et al., 2012, Campinho et al., 2014, Walter et al., 2019). Perhaps, the detailed analyses of THs content throughout development was carried out by Chang et al. (Chang et al., 2012) and Hu et al. (Hu et al., 2019) only. They revealed an increase of T_4_ before and T_3_ at the transition stage (appearance of anal and dorsal fin rays), followed by a sharp decrease of T_4_, and consistent content of T_3_ in further ontogeny.

Our data generally replicated these results, but in comparison with data represented by Hu et al. (Hu et al., 2019), the peak of T_3_ was shifted a little toward early ontogeny and occurred simultaneously with an increase of T_4_. We also observed the second increase of T_3_, which preceded the formation of the adult pigment pattern and, to our knowledge, has not been previously described. We suggest that two peaks of THs is likely a specific character of “leopard” zebrafish, and the second one is essential for the larvae-juvenile transition of skin and pigment pattern (McMenamin et al., 2014, Saunders et al., 2019, Aman et al., 2023).

The treatment with T_3_ and MPI provoked shift in THs status of experimental fish. The former increased the tissue content of T_3_, but did not affect T_4_. This result is consistent with the data reported by Walter et al. (Walter et al., 2019), and allows considering hormonal status of experimental fish as a transient T_3_ excess or acute thyrotoxicosis. In contrast, the latter caused a decrease of T_4_ and should be regarded as a T_4_ deficiency or hypothyroidism. However, we unable to detect significant changes in T_3_ level in response to MPI treatment. Similar T_3_ resilience under the methimazole and perchlorates was reported by Crane et al. (Crane et al., 2006) and Mukhi et al. (Mukhi et al., 2007). Meanwhile, at several developmental stages, we observed the developmental retardation and growth deceleration, as well as phenotypic consequences, although less pronounced but typical of hypothyroid fish (Elsalini and Rohr, 2003, Schmidt et al., 2012, Fetter et al., 2015, Salis et al., 2021, Roux et al., 2023). Taken together, these findings suggest that experimental MPI treatment caused a transient T_4_ deficiency, but effects of hypothyroidism were likely, at least in a part, mitigated by specific homeostatic mechanisms, allowing teleosts to maintain stable T_3_ level (Eales, 2019).

### Developmental effects of transient changes of THs content

As was previously shown by Keer et al. (Keer et al., 2019), the skeleton structures differed in THs sensitivity. Many, but not all, ossifications shown premature appearance and accelerated development in response to acute thyrotoxicosis, and postponed appearance and developmental retardation in response to THs deficiency. Although S. Keer with coauthors assessed the consequences of the permanently altered THs status, and we assessed the impact of transient changes, the structural responses were similar. A notable shift in the timing of appearance were detected for the ossifications composing ethmoid region and bones associated with the lateral line of the skull (frontale, sphenotic, preopercula, infraorbitals), as well as Weberian apparatus, paired fins and girdles, caudal fin rays, pharyngeal dentition and squamation. The jaw bones - dentary and maxilla, opercular series, branchiostegal rays, vertebral column, dorsal and anal fin displayed a subtle or no shift in developmental timing directly during the experimental treatment, but might demonstrate changes in the developmental rate at later stages.

In terms of the influence of THs alterations on skeleton development (Fig. 10; Supl. 6), the earliest stages (embryo and early larva) should be regarded as the weakly or less sensitive. At these stages, we failed to find changes in the growth rate and the timing of appearance of ossifications. A burst of developmental effects induced by THs alterations occurred at stages 3 and 4. Despite the growth arrest, T_3_-treated fish exhibited advanced state of the skeleton, i.e. developed more bones than euTH fish, and these bones were more massive and often had an abnormal shape due to an increased rate of ossification. Exposure to MPI led to a retardation of the overall and the skeleton development. The number and degree of ossification of the skull and postcranial skeleton were significantly less than in euTh fish.

**Fig. 10.**
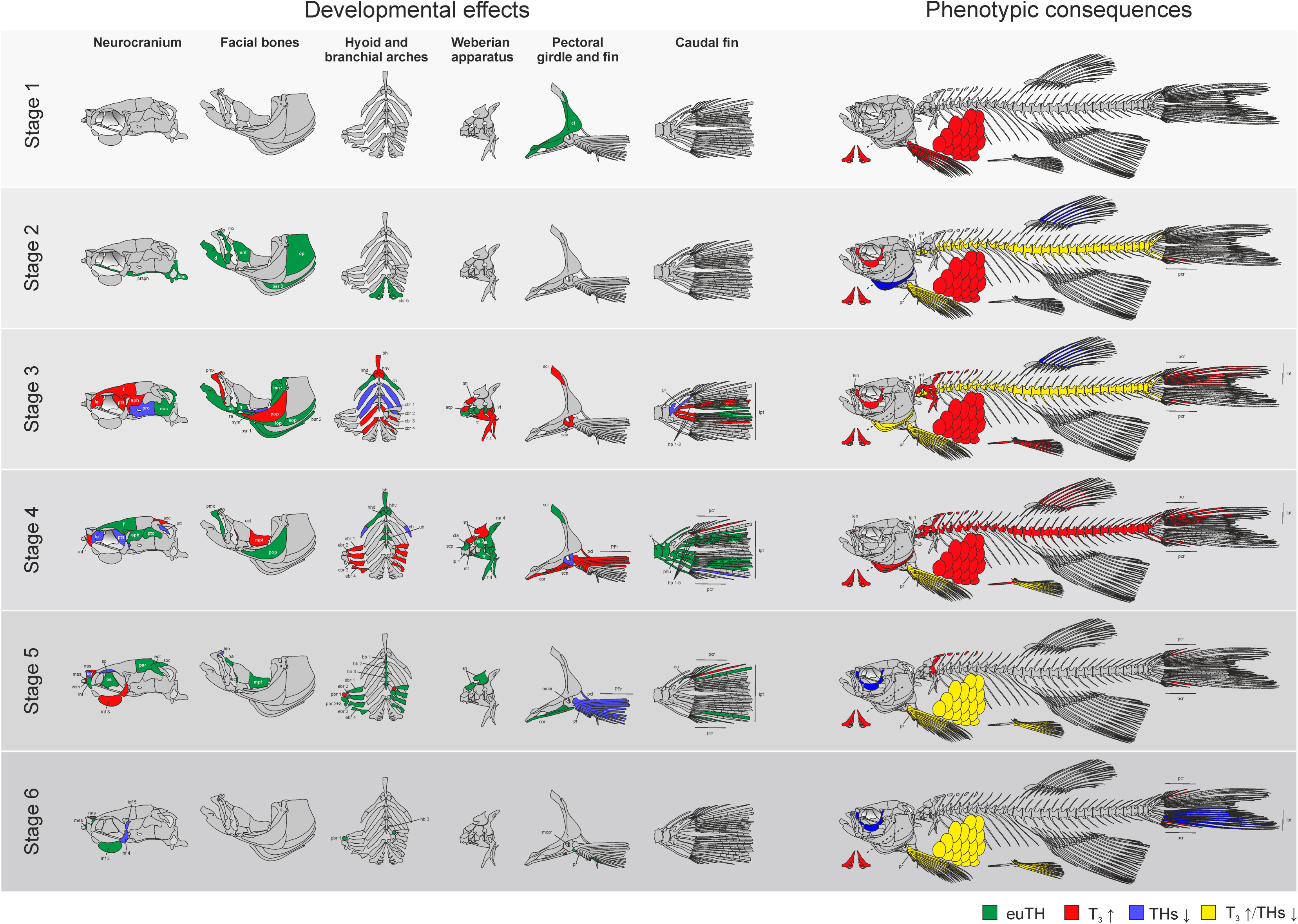
Scheme summarizing developmental effects and phenotypic consequences of transient thyrotoxicosis and hypothyroidism at different developmental stages. green – indicates timing of structure appearance in euTH fish; red indicates effects (developmental acceleration) and consequences (decrease in number, malformations) of thyrotoxicosis; blue - effects (developmental retardation) and consequences (increase in number, malformations) of hypothyroidism; yellow – phenotypic consequences (decrease or increase) of both THs excess and deficiency, respectively. Cranial ossifications: aa – anguloarticular; bb 1-3 – basibranchial (1-3); bh – basihyal; boc – basioccipital; bsr 1-3 – branchiostegal ray (1-3); cbr 1-5 – ceratobranchial (1-5); ch – ceratohyal; d – dentary; ebr 1-4 – epibranchial (1-4); ect – ectopterygoid; eh – epihyal; ent – entopterygoid; eoc – exoccipital; eot – epiotic; f – frontal; hb 3 – hypobranchial 3; hhd – dorsal hypohyal; hhv – ventral hypohyal; hm – hyomandibula; inf 1-5 – infraorbital (1-5); iop – interopercle; kin – kinethmoideum; le – lateral ethmoid; mes – mesethmoid; mpt – metapterygoid; mx – maxilla; nas – nasal; op – opercle; os – orbitosphenoid; pal – palatine; par – parietal; pbr 1 – pharyngobranchial 1; pbr 2+3 – pharyngobranchial 2+3; pmx – premaxilla; pop – preopercle; pro – prootic; ps – parasphenoid; pto – pterotic; pts – pterosphenoid; ptt – posttemporal; q – quadrate; ra – retroartucular; se – supraethmoid; so – supraorbital; soc – supraoccipital; sop – subopercle; sph – sphenotic; sym – symplectic; uh – urohyal; vom – vomer. Postcranial ossifications: cl – cleithrum; cla – claustrum; cor – coracoid; eu – epural; hp 1-5 – hypural (1-5); int – intercalarium; lp 1 – lateral process 1; lpt – lepidotrichia; mcor – mesocoracoid; na 3,4 – neural arch 3,4; pcl – postcleithrum; pcr – procurrent ray; phu – parhypural; pl – pleurostyle; pr – proximal radials; r 4 – rib 4; sca – scapula; scp – scaphium; scl – supracleithrum; sn – supraneural; tr – tripus; vt – vertebrae.

The stage 5 was also characterized by a high rate of new ossifications appearance in euTH fish. This stage might be considered as the first part of larvae-juvenile transition due to the disappearance of larval and progressive development of juvenile characteristics (Parichy et al., 2009). In general, the effects of THs alterations on the timing of bone appearance were less pronounced than at stages 3 and 4. However, T_3_ fish precociously developed traits typical of juveniles: infraorbital bones 3 and 4, mesethmoid, nasal, mesocoracoid in pectoral fin and squamation. The skull, pectoral girdle and fin were more ossified and robust, and head shape remained those of euTH fish at the end of transition period. Thus, thyrotoxicosis notably accelerated the transition to juvenile. In contrast, hypothyroidism retarded the skeleton development. At the end of the stage, MPI fish possessed the reduced number of skull bones. The MPI also caused a retardation of pectoral fin development and decrease of ossification rate in general. At the stage 6, the final part of the larva-juvenile transition, effects of hormonal treatment were far less pronounced than at previous stages and were manifested in precocious appearance of the latest skeleton structures. At the same time, an exposure with MPI caused significant retardation of growth and late skeleton structures (infraorbitals, scales) development.

### Phenotypic consequences of transient THs alterations

Stages differed in the severity of phenotypic outcomes in response to experimental manipulation with THs status (Fig. 10; Supl. 6). Treatment at the embryonic period caused an absence or weak changes in the meristic and shape characters, most of which could be neglected since they did not go beyond the variability typical of euTH fish. However, it worthy to note, that acute thyrotoxicosis led to a weak but statistically significant reduction in the scale number in fish belonging to both series, and to a decrease of the count of pharyngeal teeth and pectoral fin rays in one of two series. Meanwhile, FA level did not changed.

The effects became more noticeable at the stage 2. In response to thyrotoxicosis, both series displayed a significant decrease in the number of scales in the lateral line row, pectoral fin rays and infraorbitals, as well as the absence of intercalarium, abnormal scaphium and rib 4. Many individuals had reduced in size kinethmoideum. In series 1, the reduced number of pharyngeal teeth and vertebrae, and significant changes in head morphology were also observed. Series 2 possessed less procurrent rays in the caudal fin. MPI-treated fish were characterized by an increase in the number of rays in the pectoral and dorsal fin, branchiostegals and vertebrae. They also differed from euTH fish in head morphology.

The most pronounced consequences of experimental treatment were achieved at the stage 3 and 4. Thyrotoxicosis significantly reduced the number of the majority of meristic traits, affected the Weberian apparatus and the development of precaudal and caudal vertebrae, led to the loss of the kinethmoid, malformations and truncations in the paired and unpaired fins, including the partial or complete absence of the pelvic fin and girdle. It also influenced the overall and head morphology, and caused a surge of FA level that indicates the increase of developmental noise (Klingenberg, 2019, Zakharov et al., 2020). The results of MPI treatment were much weaker. It led to an increase in the number of several meristic traits and did not have a major effect on head morphology.

The stages 5 and 6 were characterized by a decrease of effects induced by thyrotoxicosis and an increase of the consequences of THs-deficiency. In response to hormonal treatment, only the pharyngeal dentition and number of rays in the pectoral and caudal fins were changed. MPI- treatment affected the caudal, pectoral and pelvic fin rays, the number of lateral line scales and infraorbitals. The head morphology and FA level were weakly affected by both treatments.

### Critical developmental window

The obtained results allow distinguishing critical developmental windows as for individual structures, as at organismal level. Similarly with S. Keer and coauthors (Keer et al., 2019), we observed a wide range of THs-sensitivity of the ossified skeletal structures: from subtle or no discernible reactions to a pronounced response to THs alterations. The majority of THs-sensitive bones has a defined period, when changes in THs status induces changes in the developmental timing and rate leading to phenotypic consequences, i.e. affect the phenotypic plasticity, and should be regarded as a critical developmental window (Burggren and Mueller, 2015). Some structures, for example ethmoid bones or elements of *pars auditum* composing the Weberian apparatus, are characterized by a narrow critical window, limited by a single developmental stage. Other, mostly complex traits, consisting of meristic units, like the pharyngeal dentition, squamation or pectoral fin radials and rays, have a wide critical window. Such differences in duration of the sensitive period likely associated with a multistage development of the latter. Thus, developmental pattern of pharyngeal dentition continues from early larval to transition stages and includes the development of endochondral ceratobranchial 5 and several successive tooth replacement waves (Huysseune et al., 1998) each of which is affected by THs (Woltmann et al., 2018, Keer et al., 2022). The development of squamation is a result of the prolonged and complex skin development coupled with the fish growth, and affected by THs at different stages (Aman et al., 2023, Bolotovskiy and Levin, 2015). The development of pectoral fin also undergoes several stages: larval, transitional and juvenile (Grandel and Schulte-Merker, 1998) and depends on THs (Brown, 1997, Shkil et al., 2012, Harper et al., 2023). Remarkably, that sometimes the delayed effects of THs alterations were revealed. For example, the reduction of scales in fish treated with T_3_ during the early larval stage. We suggest that in such cases, THs affect the developmental pattern of the progenitors or their environment, which, in turn, is reflected in the definitive shape of state of the structure.

Developmental stages also differ in THs-sensitivity. Since phenotypic plasticity does not significantly change in response to THs alterations at the embryonic and early larval stages, we consider these stages as the least sensitive. In contrast, the subsequent period (stages 3 and 4) is characterized by an extremely high plasticity, and should be considered a critical developmental window for skeletal development at the organismal level, even though critical windows for several structures extend beyond it. Despite the relative shortness of this period, the phenotypic consequences of acute transient thyrotoxicosis during this stage are comparable to those in fish with persistent thyrotoxicosis (Keer et al., 2019, Shkil et al., 2012, Aman et al., 2021, Keer et al., 2022).

It worthy to note, that zebrafish’s critical window to a large extend coincides with the period of increased developmental plasticity in adaptively radiating cyprinids, in particular the large African barbs *g. Labeobarbus* of Lake Tana (Shkil et al., 2015, Smirnov et al., 2012). Moreover, skeletal structures affected by THs during this period in zebrafish play the pivotal role in formation of adaptive phenotypes in various fish taxa and considered as evolutionary targets (Keer et al., 2022). These facts, along with knowledge about the role of thyroid axis in interaction between environment and genotype, as well as in adaptive radiation of various teleosts, suggest that the critical developmental window is a period of thyroid axis induced developmental decision about which developmental trajectory will be used to achieve an ecologically relevant phenotype.

## Acknowledgments

The authors would like to express their deep gratitude to Daria Kapitanova, Alexey Tsessarsky and Anna Vassilieva for their assistance in experimental works and discussion of the results. We are also grateful to Andrei Nikiforov for laboratory animals. The study was conducted in frame of the IEE RAS and IDB RAS research programs no. 0089-2021-0003 and 0088-2021-0019, respectively.

## Competing interests

The authors declare no competing or financial interests.

## Author contributions

Conceptualization: V.B., F.S.; Methodology: V.B., F.S.; Formal analysis: V.B., F.S.; Investigation: V.B., F.S.; Data curation: V.B., F.S.; Writing - original draft: V.B., F.S.; Writing - review & editing: V.B., F.S.; Visualization: V.B., F.S.

## Funding

No funding

## Data Availability Statement

All relevant data can be found within the article and the supplementary materials

## Notes

### Competing Interest Statement

The authors have declared no competing interest.

